# Do deleterious mutations promote the evolution of recombination suppression between X and Y chromosomes?

**DOI:** 10.1101/2023.11.27.568803

**Authors:** Colin Olito, Brian Charlesworth

## Abstract

An interesting new model has recently been proposed for the evolution of suppressed recombination between newly evolving X and Y chromosomes, where males are heterozygous for a locus determining sex and females are homozygous (the same principles apply to systems with female heterogamety, with the role of the two sexes reversed). The model appeals to the selective advantage that accrues to a recombination suppressor (e.g., an inversion), which arises on a male-determining haplotype that carries a smaller number of deleterious mutations than average and remains completely associated with the sex determining locus. The underlying logic of the model rests on the idea that, because such an inversion cannot become homozygous, it is “sheltered” from selection against any deleterious recessive mutations it may carry, in contrast to an autosomal inversion. It has been claimed that computer simulations of this process show that the probability that a new inversion becomes fixed within the population of Y chromosomes is substantially higher than expected under selective neutrality, and higher than that for a comparable autosomal inversion. However, analytical population genetic models of some special cases cast doubt on the magnitude of the selective advantage claimed for Y-linked inversions under this process, demanding re-evaluation of the simulation results. The published estimates of fixation probabilities of Y-linked inversions were in fact calculated after excluding inversions that were lost in the first 20 generations. This generates a substantial bias towards high fixation probabilities and obscures comparisons with neutrality. If all simulation runs are included, most parameter sets yield estimates of fixation probabilities that are close to neutral expectation, unless deleterious mutations are close to being completely recessive. The proposed sheltering mechanism is unlikely to provide a robust selective advantage to inversions suppressing recombination between evolving X and Y (or Z and W) chromosomes.

## Introduction

Understanding why, how, and when sex chromosomes stop recombining is a major goal in evolutionary biology. While a large body of population genetics research focuses on the role of sex differences in selection during the evolution of recombination supression between sex chromosomes [1, 2], several recent theoretical studies have directed attention to alternative hypotheses [3–5]. Among these alternatives is what has been called “the sheltering hypothesis” for recombination suppression between sex chromosomes [4, 6, 7].

The sheltering hypothesis postulates an initial population subject to deleterious mutation at multiple sites, resulting in a stationary probability distribution of the number of deleterious mutations per haplotype in the region of the genome spanned by a new inversion (or other recombination modifier). If genetic recombination is completely suppressed over this region in inversion heterozygotes, an inversion that arises as a unique structural mutation will be associated with a set of deleterious alleles drawn randomly from the initial probability distribution. In a randomly mating population, the inversion will experience a selective advantage if its set of alleles is associated with a higher-than-average fitness in comparison with haplotypes drawn from the initial, inversion-free population. If the inversion is sufficiently large and mutation rates are sufficiently high, the distribution of the number of mutations will be approximately normal, so that there is a 50% chance that the inversion will experience an initial selective advantage, and a 50% chance that it will be selected against. An inversion on a proto-Y chromosome (where undifferentiated and still recombining X and Y chromosomes contain the same functional sites) that completely suppresses recombination with the sex determining locus will remain completely associated with the male determining allele at this locus and can never become homozygous. In contrast, autosomal inversions can become homozygous, and may therefore experience a selective disadvantage due to homozygosity for any deleterious mutations that they carry.

Much confusion around the sheltering hypothesis arises from the mistaken assumption that the enforced heterozygosity of an inversion expanding the sex-linked region on a proto-Y chromosome also applies to the full set of alleles initially captured by the inversion. But, as was pointed out by Nei et al. [8], new mutations continue to arise at sites in inversion haplotypes that were initially free of mutations (and at all sites on non-inverted genomes), while at the same time all descendant copies of the inversion will be permanently burdened with any deleterious mutations it initially captured (ignoring back-mutation). Hence, any initial fitness advantage to the inversion will gradually decay. The best case scenario is that for initially mutation-free inversions, which will eventually become selectively neutral, while inversions initially capturing any deleterious alleles will eventually become deleterious [4, 8]. This process of fitness decay is distinct from the accumulation of deleterious mutations due to selective interference in sex-linked regions [9], and must be included in calculations concerning the ultimate fate of new inversions on both autosomes and proto-Y chromosomes [5, 10] (see [4] for a brief history of sheltering hypotheses).

An objective measure of the extent to which a selective advantage can arise in this process is the net probability of fixation of a new inversion, averaging over all possible genetic backgrounds on which a new inversion can occur, and taking into account the process of gradual decay over time of any initial selective advantage. If this probability is greater than the value for a selectively neutral mutation, there is a net selective advantage to the inversion. The fixation probability can be found either by analytical models that assume that mutations are sufficiently strongly selected against that their behaviour can be treated deterministically (i.e., *Ns* ≫ 1; [4, 5, 10], or by computer simulations of more general situations in which genetic drift as well as selection and mutation is important [3, 6, 7]).

In their recent *PLoS Biology* article, Jay et al. [6] studied the sheltering hypothesis using both deterministic and individual-based Wright-Fisher simulation models and concluded that it can explain the loss of recombination between sex chromosomes under broad parameter conditions. The authors studied several scenarios, but focused on the representative case of inversions that arise on a male-determining haplotype in a population segregating for a sex-determining locus (male heterogamety was assumed, such that males are heterozygous and females are homozygous; similar findings apply to female heterogamety). The deterministic models studied by Jay et al. [6] made the strong simplifying assumption that the initial fitness of new inversions remains constant over time. As noted above, this assumption is problematic because it ignores the spread of new mutations at loci where the inversion captured a wild-type allele. The conclusions of Jay et al. [6] therefore rest heavily on the simulation results (performed in SLiM; [11]).

## Conditional frequencies of inversion fixation

The key simulation results of Jay et al. [6] are presented in their Fig 3, and focus on populations of *N* = 1, 000 diploid individuals. Notably, their Fig 3c visualizes the proportion out of 10,000 replicate simulations where a single-copy inversion mutation either went to fixation or was segregating at a frequency of greater than 0.95 after 10,000 generations, and compares results for autosomal vs. Y-linked inversions. The result is visually striking, and suggests that autosomal inversions have little or no chance of going to fixation, while Y-linked inversions can have fixation probabilities that are many times greater than the neutral expectation of 2*/N* = 0.002. Throughout the paper, these results are interpreted and discussed as probabilities of inversion spread and/or fixation, which is not suprising; estimating fixation probabilities from the proportion of replicate simulations where fixation events occur has been common practice in theoretical population genetics for around 50 years (e.g., see [12] for an early example).

However, inspection of Jay et al. [6]’s Fig 3c raises questions. In particular, it is strange that the elevated frequencies of fixation for Y-linked inversions were especially apparent when selection was weak, even when the deleterious mutations were only partially recessive, i.e., the dominance coefficient, *h*, is only slightly smaller than 0.5 (*hs* measures the selective disadvantage to heterozygous carriers of a mutation, and *s* is the selective disadvantage to mutant homozygotes). These results are very surprising, given that *Ns* = 1 when *s* = 0.001 and *N* = 1, 000, so that the deleterious mutations are nearly neutral with respect to their population dynamics [13]. Intuitively, it is difficult to understand how nearly neutral mutations could generate such a large selective advantage to a new inversion.

Fortunately, standard neutral population genetics theory can be used to provide an upper bound to the expected fixation probability of a new inversion in this case, as described in Appendix A of Online Supplement S1. Specifically, with weak selection (*Ns* ≤ 1) against deleterious mutations, the approximate fixation probability for a new single-copy Y-linked inversion relative to the neutral value of 2*/N* can be expressed as:

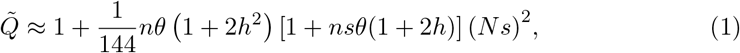

where *n* is the number of sites spanned by the inversion and *θ* = 4*Nμ* (ignoring second order terms in the product of *N* and the selection coefficient). The approximation works well for larger *h*, but increasingly underestimates inversion fixation probabilities for partially to completely recessive mutations. For the parameter values just mentioned, and assuming *h* = 0.4, this upper-bound for the fixation probability is 1.074 times the neutral value (corresponding to an absolute survival probability of 0.00215) for *μ* = 10^−9^ (where *μ* is the per-site mutation rate), and 1.839 times the neutral value for *μ* = 10^−8^ (with absolute survival probability of 0.00368), far smaller than the estimates of Jay et al. [6]. Indeed, one can calculate several analytic approximations for the upper-bound of possible inversion fixation probabilities in both strong and weak selection scenarios, none of which approach the reported values (see Appendix A).

The resolution of this paradox can be found in a brief statement in the caption to Fig 3 of Jay et al. [6], where they state that “*Only inversions not lost after 20 generations are considered here*”. Identical statements appear in the captions of all supplementary figures showing fractions of fixed inversions. Inspection of Jay et al. [6]’s computer code used to generate the original figure confirms that inversions going extinct in the first 20 generations were excluded from the dataset before calculating and plotting the fractions of fixed inversions (see S1: Appendix B; initially mutation-free inversions were also excluded, with minimal effect). Hence, what Jay et al. [6] actually plotted were frequencies of inversion fixation conditioned on their early survival. To put the effect of this condition into context, a simple approximation due to Fisher [13, pp.80–85] for the cumulative probability of gene extinction over time suggests that Jay et al. [6] were excluding around 90% of the extinctions occuring in their simulations prior to calculating the frequency of fixed inversions. Hence, ignoring inversions lost within 20 generations will greatly overestimate the true fixation probabilities.

Further examination of Jay et al. [6]’s supplementary computer code reveals that they also performed simulations where segregating mutations were selectively neutral (*s* = 0.0; and therefore inversions were also selectively neutral), but these data were excluded from all figures, and were never discussed in the article, analogous to an experimental study omitting any presentation or discussion of results for a control treatment. Including the data for neutral inversions underscores the conditional nature of Jay et al. [6]’s results, as neutral inversions appear to go to fixation at frequencies many times greater than the theoretical expectation of 2*/N* and very close to the observed frequencies for *s* = 0.001 (when *μ* = 10^−9^; see Fig 1). Plotting the data for neutral inversions also highlights the fact that, for the larger mutation rate (*μ* = 10^−9^), the fractions of fixed Y-linked inversions drop below the neutral results as mutations become less recessive (as *h* approaches 0.5).

**Fig 1.**
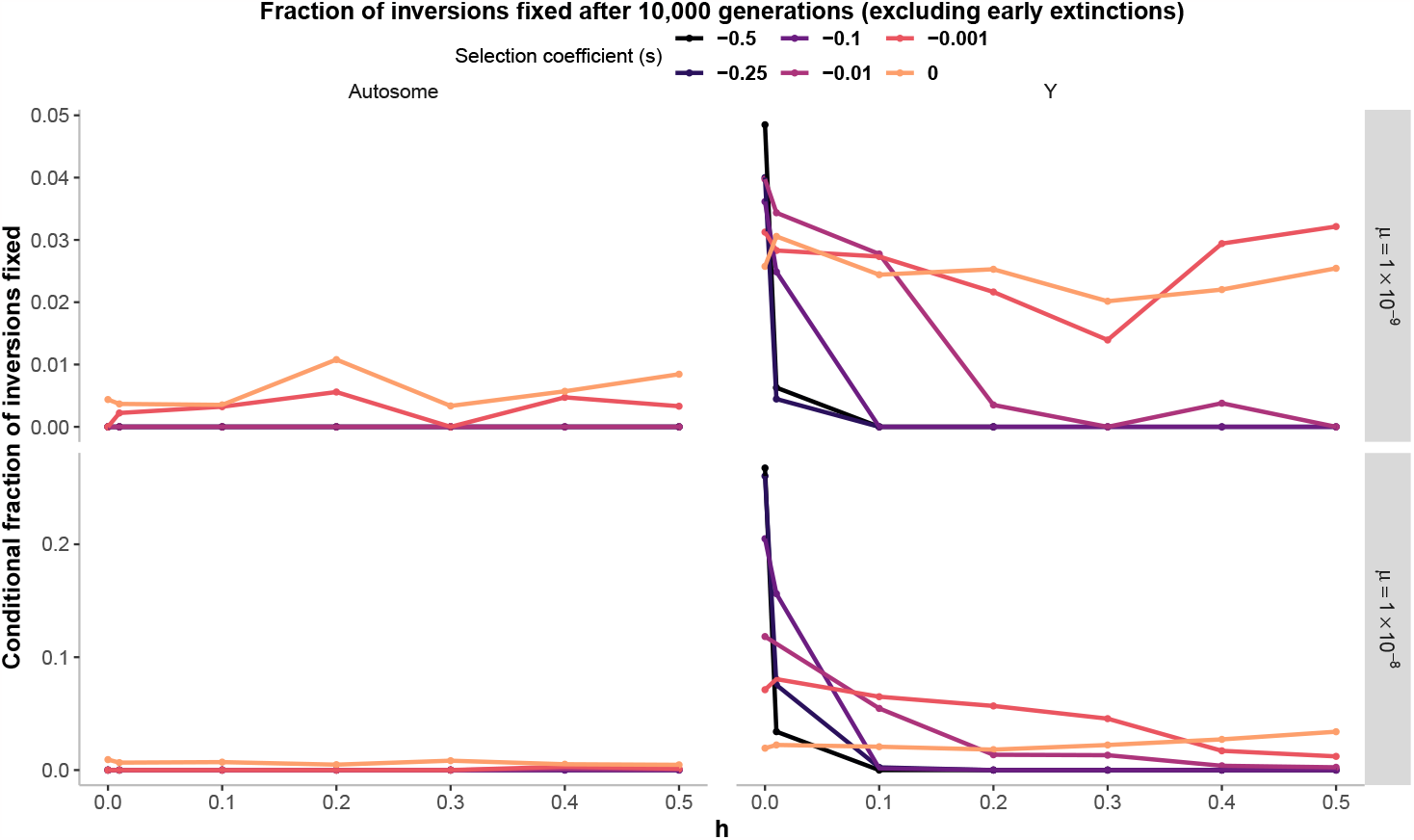
Corrected presentation of **Fig 3c** from Jay et al. [6]. We have included the data for neutral mutations (where *s* = 0.0), and modified the y-axis label and main figure title to accurately reflect the correct interpretation of the data. Specifically, the figure displays the frequency among 10,000 replicate simulations where independent single-copy inversion mutations go to fixation *conditioned on those inversions surviving longer than 20 generations, and initially capturing at least one mutation*. Left-hand panels show results for autosomal inversions, while right-hand panels show results for inversions expanding the sex-linked region on a proto-Y chromosome. Results are shown for a population size of *N* = 1, 000, dominance coefficients varying between completely recessive to additive fitness effects (*h* = *{*0.0, 0.01, 0.1, 0.2, 0.3, 0.4, 0.5*}*), selection coefficients ranging from quite weak to extremely strong selection (*s* = *{*0.0, 0.001, 0.01, 0.1, 0.25, 0.5*}*), and two per-base-pair mutation rates (*μ* = *{*10^*−*9^, 10^*−*8^*}*).

*We emphasize that by excluding data for neutral inversions and plotting inversion fixation frequencies conditioned on their early survival, Jay et al. [6] presented a subset of their data that was not appropriate for addressing their biological question, and which obscured comparisons between their data and neutral fixation probabilities*. As we show below, calculating inversion fixation frequencies using Jay et al. [6]’s full dataset sheds a very different light on the extent to which their simulation results support their sheltering hypothesis.

## Re-interpreting Jay et al. (2022)’s simulation results

Calculating and plotting the frequencies of inversion fixation using Jay et al. [6]’s full dataset (Fig 2; see also S1: Appendix C, Supplementary Figs S1–S8) immediately demands a reinterpretation of their results. There are several important conclusions.

**Fig 2.**
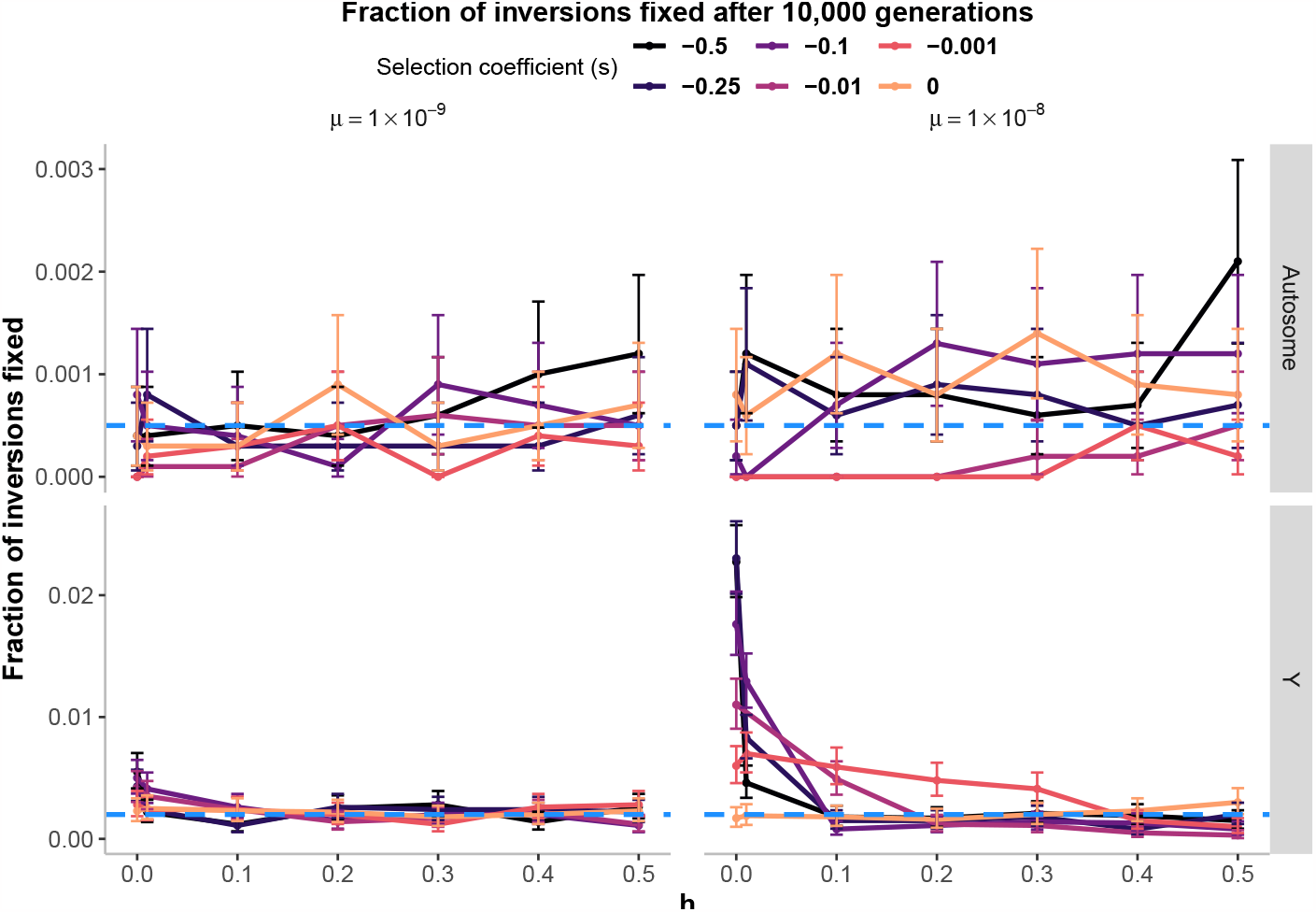
A revised presentation of the results from Jay et al. [6]’s **Fig 3c**, where the fraction of fixed inversions has been calculated using the full dataset, the figure panels have been re-arranged so that autosomal and Y-linked inversions are each plotted on their own y-axis scale, and Poisson 95% confidence intervals have been added to each estimate. These results can be interpreted as estimates of the overall probability of fixation for the inversions. Results are shown for a population size of *N* = 1, 000, dominance coefficients varying between completely recessive to additive fitness effects (*h* = {0.0, 0.01, 0.1, 0.2, 0.3, 0.4, 0.5} ), selection coefficients ranging from quite weak to extremely strong selection (*s* ={ 0.0, 0.001, 0.01, 0.1, 0.25, 0.5007D ), and two per-base-pair mutation rates (*μ* = {10^*−*9^, 10^*−*8^ }). The expected proportion of fixations for a neutral gene first arising as a single copy is indicated by the blue horizontal dashed line (corresponding to 1*/*(2*N* ) for an autosomal gene, and 2*/N* for a Y-linked gene).

First and most obvious, the frequencies of inversion fixations are generally close to neutral expectations, with the notable exception of Y-linked inversions capturing many loci segregating for nearly completely recessive deleterious mutations (*h* ≤ 0.01; but see below). The results for Y-linked inversions with weak selection and the higher mutation rate (*s* = 0.001, *h >* 0.25, and *μ* = 10^−8^) are also reasonably well approximated by Eq (1). However, the frequency of fixation for Y-linked inversions appears to drop below 2*/N* and approach zero for *Nhs >* 1 when *μ* = 10^−8^. Nevertheless, all of the simulation results for *N* = 1, 000 must be interpreted with caution, as even the autosomal inversions, which have a larger effective population size, exhibit nearly neutral dynamics. It should be noted that Fig 3a of Jay et al. [6] presents one of the minority cases where no autosomal inversions went to fixation; in fact some autosomal inversions went to fixation in 58 out of the 70 parameter sets explored in their Fig 3c.

Second, elevated fixation frequencies of Y-linked inversions when deleterious mutations are completely recessive or nearly so are also seen in the full dataset. In this case, a similar approximation for the upper bound of the expected fixation probability to that presented in Eq (1) is also possible (see S1: Appendix A.2.2). When deleterious mutations are completely recessive and selection is strong (*Ns* ≫ 1), the approximate relative fixation probability of a Y-linked inversion can be expressed as:

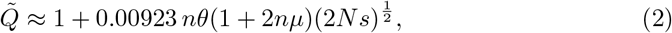

and the simulation results presented in Fig 2 generally agree well with this approximation (see S1: Appendix A2.2). Importantly, Eq (2) predicts that the fixation probability for a Y-linked inversion subject to completely recessive mutations increases linearly with the mutation rate and with the square root of 2*Ns*. This effect is clearly visible in Fig 2 and in S1 Appendix C: Fig S2–S5 (corresponding to Figs S15–S16 of Jay et al. [6]).

The plausibility of the completely recessive selection scenario deserves comment, however. Current empirical data and theory suggests an average dominance coefficient for mildly deleterious mutations of *h* ≈ 0.25 [14–17]. The inversions modelled by Jay et al. [6] are also very large: capturing from 500 kb to 5 Mb of *selected sites*. Except in species with compact genomes, the large amount of nearly neutral intergenic and intronic sequences mean that this would correspond to truly massive inversions, and an unrealistically high incidence of completely recessive deleterious mutations across the genome [14].

A related issue is that Jay et al. [6] simulated small populations, but used standard per-site mutation rates, resulting in very small population scaled mutation rates, even compared with humans, for which silent nucleotide site diversity is approximately 10^−3^ [18] (as compared with a value of 4 × 10^−5^ for a mutation rate of 10^−8^ with *N* = 1, 000). On the other hand, they simulated inversions with many selected sites. Assuming that the product of the number of sites and the mutation rate is what matters for the fate of an inversion, this roughly balances out as far as biological realism is concerned. However, the results are unlikely to scale well when using different mutation rates or inversion sizes, so that the question of how inversion size influences fixation probability is not adequately addressed by these simulations.

Lastly, in their supplementary material, Jay et al. [6] present selected results for a larger population size (*N* = 10, 000, with *μ* = 10^−8^) which we have also replotted to show correct inversion fixation frequencies (see Supplementary Figs. S4–S6). Reassuringly, fixation of autosomal inversions becomes exceedingly rare in larger populations, as predicted by prior theory focusing on large populations [8, 10]. The elevated Y-linked inversion fixation frequencies for values of *h* ≤ 0.01 appear robust to population size. However, for partially recessive mutations (where *h >* 0.01) the results are still mostly indistinguishable from neutrality and the precision of the estimates (based on 10,000 replicate simulations) is too low to reliably estimate such small fixation probabilities (see Supplementary Fig S6).

## Concluding remarks

By omitting data for neutral inversions and then plotting conditional inversion fixation frequencies and incorrectly labeling and interpreting them as estimates of overall inversion fixation probabilities, Jay et al. [6] presented a misleading picture of how well the simulation data supported their hypothesis. When using their full dataset to calculate inversion fixation frequencies, it becomes clear that their model predictions are indistinguishable from neutral dynamics for most for the parameter conditions studied. Our new analytic approximations of the upper bound of inversion fixation probabilities confirm that the corrected results are consistent with classical population genetics theory, and inform our intuition about how these inversions are expected to behave under broader parameter conditions than those simulated. Taking into consideration the corrected simulation results, along with our new analytic approximations, it seems that the frequency of fixation for new inversions expanding the sex-linked region on a proto-Y chromosome is unlikely to differ much from neutral expectations, unless the mutations in question are nearly completely recessive with respect to their effects on fitness. Indeed, for many biologically plausible parameter sets, the frequencies will be lower than neutral. The sheltering mechanism cannot, therefore, be regarded as robustly providing a selective advantage to new inversions. This does not, of course, mean that inversions evolving under indirect selection due to segregating deleterious mutations cannot go to fixation on evolving Y chromosomes; it simply means that their rate of establishment is likely to be close to, and often lower than, that expected under selective neutrality.

## Data Availability Statement

The original computer code and data for Jay et al. (2022) is available both as an online supplement to this article, and is also archived on Zenodo (https://zenodo.org/records/10041542). Computer code needed to reproduce the figures in this article and the SI are also available as an online supplement, and on GitHub (https://github.com/colin-olito/Jay-etal-Comment). An official version of record at the time of submission is archived on Zenodo (https://zenodo.org/record/8417368,doi: 10.5281/zenodo.8417368).

## Supporting information

**S1. Supplementary Appendices A–C**. Appendix A: Approximate upper bounds for inversion fixation probability; Appendix B: Jay et al. (2022)’s Computer Code; Appendix C: Supplementary Figures.

**S2. Computer code to reproduce our results**. R code to reproduce the key simulation figures from Jay et al. (2022), as well as our reanalysis of the full datasets and modified versions of their figures presenting correcly-calculated inversion fixation frequencies.

**S3. Jay et al. (2022)’s Original Computer Code** Unaltered code to run the analyses presented in Jay et al. (2022).

## Acknowledgements

CO is supported by a VR Establishment grant (no. 2022-03603). Discussions with Deborah Charlesworth, Chandra Venables, Tim Connallon, Crispin Y. Jordan, and P. Czuppon greatly improved the manuscript. The authors have no conflicts of interest to declare.

## Notes

https://zenodo.org/records/8417368

### Competing Interest Statement

The authors have declared no competing interest.

