## Appendix for "Do deleterious mutations promote the evolution of recombination suppression between X and Y chromosomes?"

### Supporting Information for: Do deleterious mutations promote the evolution of recombination suppression between X and Y chromosomes?

Colin Olito[1,*] and Brian Charlesworth[2]

November 27, 2023

[1] Department of Biology, Lund University, Lund 223 62, Sweden;
[2] Institute of Ecology and Evolution, School of Biological Sciences, University of Edinburgh, Edinburgh EH9 3FL, UK;

Computer code needed to reproduce the simulations and main figures is available on GitHub (`https://github.com/colin-olito/Jay-etal-2022-Comment`).

**Appendix B Jay et al. (2022)'s Computer Code** **9**

**Appendix C Supplementary Figures** **12**

### Appendix A Approximate upper bounds for inversion fixation probability

For all of the population genetic models considered here, following Jay et al. (2022), we start with the initial condition of a proto-Y chromosome in a diploid population in which the X and Y are completely undifferentiated (i.e., with the same functional sites and recombination between them). We also assume no sex differences in mutation rates or selection coefficients. Under these assumptions, one can derive a number of analytic approximations for the expected fixation probability of an otherwise neutral chromosomal inversion (i.e., neutral in the absence of standing genetic variation in the region in question) that expands the sex-linked region on a proto-Y chromosome in the presence of deleterious mutational variation. The simplest of these approximations make the same simplifying assumption that Jay et al. (2022) made in their deterministic models: that the initial relative fitness of the inversion will remain constant over time (i.e., ignoring time-dependent selection due to recurrent deleterious mutations). Approximations made under this assumption therefore represent upper bounds for the possible probabilities of inversion fixation. The key step is then to approximate the expected initial relative fitness of an inverted Y relative to standard-arrangement Y chromosomes, averaging over all of the possible genetic backrounds in which the inversion may arise. There are two main approaches one can take which correspond roughly to strong- vs. weak-selection scenarios (i.e., high- vs. low-$Ns$). For weak-selection scenarios, established results from standard neutral theory can be used to simplify the approximations. For strong-selection scenarios, one can simplify the mathematics considerably by assuming that deleterious mutations are present at mutation-selection balance. We focus on the case of weak selection below, which is central to understanding why the large fixation probabilities found by Jay et al. (2022) for weak selection scenarios were so peculiar. We then turn our attention to approximations for the case of strong selection (i.e., $Ns > 1$). Throughout the appendix, we focus on parameter values used in Figure 3c of Jay et al. (2022), where $N = 1{,}000$, $\mu = \{10^{-9}, 10^{-8}\}$, and we denote the number of selected sites spanned by the inversion by $n = 2 \times 10^6$. Other parameters are the same as described in the main text.

#### A.1 Approximations for weak selection

In developing analytical approximations for the case of weak selection, it is useful to note that the default setting for the SLiM simulation package used by Jay et al. (2022) only tracks segregating loci. If a deleterious mutation goes to fixation, the event is counted as a substitution, and the newly fixed mutation is immediately treated as a new wild-type allele. For nearly neutral mutations with a low per nucleotide site mutation rate, this behaviour means that SLiM simulates the infinite sites model, according to which the mutation process is unidirectional and only a single mutation segregates in the population at a given site during its sojourn in the population (Fisher 1930; Kimura 1971). In this case, we can use predictions from infinite sites neutral theory (Watterson 1975) to verify that the simulations are behaving as one would expect. Under strict neutrality, the expected number of segregating sites in a sequence of $n$ basepairs for an autosomal locus in a Wright-Fisher population of size $N$ is equal to:

$$n_s = n\theta \left[\ln(2N) + 0.5772\right], \tag{A1}$$

where $\theta = 4N\mu$. With a large number of sites, as in the Jay et al. (2022) simulations, the observed number of segregating sites should be close to $n_s$.

The neutral p.d.f. for allele frequency $q$ is $\theta q^{-1}$ (Kimura 1971; Ewens 2004); integrating the product of this with $q$ between $q = 1/2N$ and $1 - 1/2N$, which is close to integration between 0 and 1 with large $N$, yields the expected number of mutations per haploid genome over all segregating sites, $\bar{n} \approx n\theta$. The mean frequency of a mutation at a segregating site is thus:

$$\overline{q} = \frac{\overline{n}}{n_s} = \frac{1}{\ln(2N) + 0.5772} \tag{A2}$$

These predictions can be compared with the results in Figure S25 of Jay et al. (2022) for the nearly neutral case of $s = 0.001$, $N = 1000$, $n = 2 \times 10^6$, and $\mu = 10^{-8}$. Substituting these numbers into Equations (A1) and (A2), we obtain $n_s = 6,542$ and $\overline{q} = 0.122$; these agree closely with the equilibrium results in Figures S25a and S25b for low $h$ values, which are most likely to behave as nearly neutral. For the higher $h$ and $s$ values shown in their Figure S25, the neutral predictions overestimate $n_s$ and $\overline{q}$.

We can therefore be safe in assuming that neutral theory will provide upper bounds on $n_s$ and $\overline{q}$ for the case of weak selection ($Ns \approx 1$). We can obtain an upper bound on the fixation probability of a new inversion by ignoring the process of decay of any initial advantage. For a site with frequency $q$ of a deleterious mutation, there is a probability $p = 1 - q$ that a random inversion haplotype carries a wild-type allele, and a probability $q$ that it carries a mutant allele. In the first case, its fitness relative to wild-type is $w_1 = 1 - qhs$; in the second case, it is $w_2 = 1 - s(ph + q) = 1 - s[h + q(1-h)]$. The mean fitness of the population at this site, conditional on $q$, is $\overline{w} = 1 - sq[2h + q(1-2h)]$. The deviations of the $w_i$ from $\overline{w}$ are as follows:

$$\delta w_1 = sq[h + q(1-2h)] \tag{A3a}$$

$$\delta w_2 = -s\left[h + q(1-3h) - q^2(1-2h)\right] \tag{A3b}$$

It is easily verified that the mean fitness deviation, $p\delta w_1 + q\delta w_2$, is equal to zero. If the $\delta w_i$ for each site are small, and all sites have the same mutation rate and selection coefficient, the variance in the fitness deviation of an inversion under a multiplicative or additive fitness model (assuming that the sites evolve independently of each other) is given by:

$$V_{\delta w} \approx n_s \int_0^1 \left[p\delta w_1^2 + q\delta w_2^2\right] \phi_s(q)\, dq \tag{A4}$$

where $\phi_s(q)$ is the probability density of $q$ for a segregating site.

To obtain the variance in the selective advantage of a new inversion, $V_{\delta w}$ should be normalised by the square of the mean fitness with respect to all segregating sites, $\overline{w}_s$. With additive fitness effects, we have:

$$\overline{w}_s = 1 - n_s s \int_0^1 q[2h + q(1-2h)]\, \phi_s(q)\, dq. \tag{A5}$$

If this is expression is close to 1, it provides a good approximation to the multiplicative case; otherwise, the following expression should be used:

$$\overline{w}_s = \exp\left\{-n_s s \int_0^1 q[2h + q(1-2h)]\phi_s(s)\, dq\right\} \tag{A6}$$

Provided that there is independence among sites, these expressions are in fact completely general under the specified assumptions. To obtain useful results, we can use the neutral approximation for weak selection:

$$n_s \phi_s(q) = \frac{n\theta}{q}. \tag{A7}$$

Inserting this expression into Equation (A4), after considerable algebra we obtain:

$$V_{\delta w} \approx \frac{1}{12} n\theta(1 + 2h^2)s^2. \tag{A8}$$

We also have

$$n_s s \int_0^1 q[2h + q(1-2h)]\, \phi_s(q)\, dq = \frac{1}{2} n\theta(1 + 2h)s. \tag{A9}$$

If this quantity is sufficiently small, as is the case for the parameter values described above, we obtain the following expression for the variance in the selection coefficient of a new inversion:

$$V_{si} \approx \frac{1}{12} n\theta(1+2h^2)\left[1+n\theta s(1+2h)\right]s^2. \tag{A10}$$

These results can be used to obtain an expression for the net fixation probability of a new inversion. Assuming a 1 : 1 sex ratio, the population size for the Y chromosome subpopulation is $N/2$, so that the neutral fixation probability of a new mutation is $2/N$. Let $\gamma = N\delta w/\overline{w}_s$ be the population scaled selection coefficient for a Y-linked inversion. A slight modification of the standard diffusion approximation for the fixation probability for an autosomal mutation (Fisher 1930; Kimura 1962) to fit the case of Y-linkage gives the fixation probability relative to the neutral value as:

$$\tilde{Q} \approx \frac{\gamma}{1-e^{-\gamma}} \tag{A11}$$

The right hand side of Eq (A11) is also the exponential generating function for the Seki-Bernoulli numbers:

$$\frac{x}{1-e^{-x}} = \frac{x}{2}\left(\coth\frac{x}{2}+1\right) = \sum_{m=0}^{\infty}\frac{B_m^+ x^m}{m!}. \tag{A12}$$

This means that our expression, Eq (A11), can be expressed as a Taylor series expansion where the coefficients of $1/m!$ are the Seki-Bernoulli numbers, $B_i^+$. $B_0^+ = 1$ and $B_1^+ = \frac{1}{2}$ is the only non-zero $B_i^+$ with an odd numbered subscript, such that $B_2^+ = \frac{1}{6}$, $B_4^+ = -\frac{1}{30}$, $B_6^+ = \frac{1}{42}$, $B_8^+ = -\frac{1}{30}$, etc. We can therefore approximate Eq (A11) to arbitrary precision using:

$$\tilde{Q} \approx 1 + \mathbb{E}\left\{\frac{1}{2}\gamma + \frac{1}{12}\gamma^2 - \frac{1}{720}\gamma^4 + \frac{1}{30240}\gamma^6 + \dots\right\}. \tag{A13}$$

The expectation of $\gamma$ is 0. For small $\gamma$ values, as in the present case, only the term in $\gamma^2$ needs to be considered, so the probability of fixation relative to neutrality can be written as:

$$\tilde{Q}_2 \approx 1 + \frac{1}{12}V(\gamma) \tag{A14a}$$

$$\approx 1 + \frac{1}{144} n\theta\left(1+2h^2\right)\left[1+ns\theta(1+2h)\right](Ns)^2 \tag{A14b}$$

With $n = 2\times10^6$, $\mu = 10^{-9}$, $N = 1{,}000$, $h = 0.4$ and $s = 0.001$ ($Ns = 1$), the relative survival probability is 1.074 and the absolute survival probability is 1/500 times this, i.e., 0.00215. With $\mu = 10^{-8}$, the corresponding values are 1.839 and 0.00368, respectively. We reiterate, however, that this represents an upper bound for the inversion fixation probabilities, as it ignores the subsequent decay of any initial fitness advantage for inversion haplotypes capturing fewer than the average number of deleterious alleles.

#### A.2 Approximations for strong selection

For strong-selection secenarios ($Ns \gg 1$), analogous approximations for the upper bound of inversion fixation probabilities essentially represent a best-case scenario under Jay et al. (2022)'s proposed sheltering mechanism, which emphasized the role of selection in driving inversion fixations. Below, we derive simple approximations under the key assumption that the population is initially at equilibrium under mutation and selection balance when deleterious mutations are recessive or partially recessive. We first deal the more general case where deleterious mutations are partially recessive ($0 < h < 0.5$), then conclude by dealing with the case of completely recessive deleterious mutations ($h = 0$), which corresponds to the highest fixation probabilities for Y-linked inversions in Jay et al. (2022)'s simulation results.

##### A.2.1 Partially recessive deleterious mutations

An initially mutation-free inversion will have the maximum possible initial relative fitness, and hence will also have the highest probability of fixation. Assuming weak selection and mutation, the marginal fitness of a standard Y chromosome will be $w_Y \approx e^{-2U}$ (the same as an autosomal inversion of the same size), and that of a new mutation-free inverted Y will be $w_I \approx e^{-U}$, where $U = n\mu$ is the mutation rate over the genomic region spanned by the inversion. The relative fitness of the new mutation-free inversion is thus $w_I/w_Y = e^U \approx 1+U$, with the approximation working well when $U$ is small, as it is for the parameter values considered by Jay et al. (2022) and on which we focus below. For the mutation rates given above, we have:

$$\frac{w_I}{w_Y} \approx 1+n\mu = 1+(2\times 10^6 \times 10^{-9}) = 1.002 \tag{A15a}$$

$$\frac{w_I}{w_Y} \approx 1+n\mu = 1+(2\times 10^6 \times 10^{-8}) = 1.02 \tag{A15b}$$

With the selective advantage, $U$, and using the diffusion approximation for the probability of fixation (Fisher 1930; Kimura 1962), we have

$$\Pr(\text{fix}) \approx \frac{2U}{1-e^{-NU}} = 0.004 \tag{A16}$$

for $\mu = 10^{-9}$. For $\mu = 10^{-8}$, we get a maximum fixation probability of $\approx 0.04$.

These simple approximations highlight the fact that, even under the most optimistic interpretation of the selection process proposed by Jay et al. (2022) (i.e., selection is strong, all inversions are initially mutation-free, and there is no decay of an inversion's initial fitness benefit over time), Y-inversion fixation probabilities are expected to be on the order of $U$, the overall mutation rate over the chromosomal region in question. For the parameter conditions emphasized in Fig 3c of Jay et al. (2022), this upper bound is around twice the neutral value of $2/N$ for $\mu = 10^{-9}$. As noted in the main text, however, the actual fixation probabilities for strong-selection scenarios will generally be much lower due to the time-dependent selection process experienced by these inversions, and the fact that only a fraction of inversions will be initially mutation free.

##### A.2.2 Completely recessive deleterious mutations

Here, we consider the case of completely recessive deleterious mutations with $Ns \gg 1$, which the simulation results of Jay et al. (2022) show is the most favourable case for the establishment of inversions that expand the sex-linked region on a Y chromosome. In this case, the process of fitness decay during the spread of a Y-linked inversion is minimal, as any new mutations that it carries are not expressed unless they are paired with a corresponding mutation on an X chromosome (which happens very rarely). Population genetics theory has shown, however, that drift can cause large departures from mutation-selection balance when mutations are recessive or close to recessive (Wright 1937; Nei 1968), so that we need to consider the effects of drift in our analysis.

Nei (1968) analysed the properties of the stationary distribution of $q$ with selection against completely recessive deleterious alleles. A true stationary distribution requires mutations to occur in both directions, whereas the SLiM simulations of Jay et al. (2022) involve mutations only from wild-type alleles to deleterious alleles. However, with large $Ns$ the proportion of sites fixed for deleterious mutations is negligible, so that the properties of this distribution are very similar to the case when there is one-way mutation from wild-type to mutant alleles, which involve a more complex expression (see Equation 9.20 of Kimura 1964). The form of this equation is such that it imposes a more severe penalty on large values of $q$ than the stationary distribution, so that the following calculations will somewhat overestimate the mean value of $q$ at segregating sites, although the effect is unlikely to be large when $Ns \gg 1$.

Nei (1968) (see also Wright 1937) showed that the unconditional mean allele frequency when $2N\mu \ll 1$ (as in the present case) is given by:

$$\overline{q}_u \approx \mu\sqrt{2\pi N/s}. \tag{A17}$$

The probability that a site is fixed for the wild-type allele is given by:

$$P_f \approx \int_0^{1/2N} \phi(q)dq \tag{A18}$$

where $\phi(q)$ is the unconditional p.d.f. for $q$. The probability that a site is fixed for the mutant allele is negligible when $Ns \gg 1$, so that the probability that a site segregates for mutant and wild-type alleles is $P_s = 1 - P_f$.

For further progress, it is convenient to use the transformation $t = q^2$. As shown by Nei (1968), the p.d.f. for $t$ is a gamma distribution, provided that $Ns \gg 1$:

$$\begin{aligned}\phi(q) &\approx \Gamma\left(\frac{\theta}{2}\right)^{-1} S^{\left(\frac{\theta}{2}\right)} \exp(-St) t^{\left(\frac{\theta}{2}-1\right)} \\ &\approx \frac{\theta}{2} S^{\left(\frac{\theta}{2}\right)} \exp(-St) t^{\left(\frac{\theta}{2}-1\right)},\end{aligned} \tag{A19}$$

where $S = 2Ns$.

From Equation (15) of Nei (1968), the unconditional expectation of $q^2$ is equal to $V_{qu} + \overline{q}_u^2 = \mu/s$, where $V_{qu}$ is the variance of $q$ over its unconditional distribution. Taking the product $n_s = nP_s$ and multiplying by the conditional expectation of $q^2$, $\mu/P_s s$, we obtain:

$$\overline{w}_s \approx 1 - n\mu. \tag{A20}$$

This result is identical to the value under mutation-selection balance.

We can now proceed on the same lines as for the neutral case. We can safely assume that $q \ll 1$, so that Equations (A3), with $h = 0$, can be written as follows:

$$\delta w_1 = sq^2 \tag{A21a}$$

$$\delta w_2 \approx -sq \tag{A21b}$$

As shown above, when considering the Y subpopulation, the scaling of the selection coefficient by population size involves $Ns$ not $2Ns$. In order to use the equivalent of Eq (A13) we need to obtain the relevant moments of $\delta w_i$. The expectation of the scaled values of the $\delta w_i^j$ is given by:

$$\left(\frac{S}{2}\right)^j \int_0^1 \left\{(1-q)q^{j+2} + q^{j+1}\right\} \phi(q)\,dq \approx \left(\frac{S}{2}\right)^j \int_0^1 q^{j+1}\phi(q)\,dq. \tag{A22}$$

Using the transformation $t = q^2$, we can make use of the fact that for $\phi(q)$, which is gamma distributed, we have:

$$\begin{aligned}q^{j+1}\phi(q) &= t^{(\theta+j+1)/2}\phi(t) \\ &= \Gamma\left(\frac{\theta}{2}\right)^{-1} S^{\frac{\theta}{2}} e^{-St} t^{\frac{\theta+j}{2}-1}\end{aligned} \tag{A23}$$

As in Eq (A13), only even values of $j$ are of interest. The $(j+1)th$ moment of $\phi(q)$ is obtained by integrating this quantity between 0 and 1, and is proportional to a gamma distribution with shape parameter $(\theta + j + 1)/2$ and scale parameter $S$, except that the normalization constants $(\Gamma(\theta+j+1)/2)^{-1}$ and $S^{(\theta+j+1)/2}$ appropriate for this distribution are replaced with $\Gamma\left(\frac{\theta}{2}\right)^{-1} \approx \frac{\theta}{2}$ and $S^{\frac{\theta}{2}}$, respectively. The resulting integral is approximately equal to:

$$\begin{aligned}\int_0^1 q^{j+1}\phi(q)\,dq &\approx \frac{\theta}{2}\Gamma\left(\frac{\theta+j+1}{2}\right)S^{-\frac{j+1}{2}} \\ &\approx \frac{\theta}{2}\Gamma\left(\frac{j-1}{2}+1\right)S^{-\frac{j+1}{2}}\end{aligned} \tag{A24}$$

Using the relation $\Gamma(x+1) = x\Gamma(x)$ (for $x > 0$), this simplifies to:

$$\frac{\theta}{2}\frac{j-1}{2^{j+2}}\Gamma\left(\frac{j-1}{2}\right)S^{-\frac{j+1}{2}} \text{ for } (j = 2, 4, 6, 8, \ldots). \tag{A25}$$

Using Eq (A22) and normalising by the population mean fitness, we obtain the following expression for the expectation of the *jth* power of the scaled selection coefficient $\gamma$, which can be substituted into Eq (A13):

$$\mathbb{E}\left\{\gamma^j\right\} \approx \frac{\theta(j-1)}{2^{j+2}}\Gamma\left(\frac{j-1}{2}\right)S^{\frac{j-1}{2}}(1-2n\mu)^{-j}. \tag{A26}$$

Using only the term in $j = 2$, and using $\Gamma(0.5) = 1.773$, we obtain the equivalent of Eq (A11) for the net fixation probability of a new inversion relative to the neutral value with strong selection against completely recessive deleterious mutations:

$$\tilde{Q} \approx 1 + \frac{1}{12}V_\gamma \approx 1 + 0.00923\,n\theta(1+2n\mu)S^{\frac{1}{2}}. \tag{A27}$$

With $n = 2\times10^6$, $\mu = 10^{-9}$, $N = 1,000$, and $s = 0.01$ ($S = 20$), we have $\theta = 4\times10^{-6}$ and $n\theta = 8$, so that the relative survival probability is 1.332 and the absolute survival probability is 1/500 times this, i.e., 0.00266. With $\mu = 10^{-8}$, the corresponding values are $\theta = 4\times10^{-5}$, and $n\theta = 80$, so that the relative fixation probability is 4.434 and the absolute fixation probability is 0.00886. These numbers are all very close to the simulation values in Fig 2 of the main text. An important conclusion is that the fixation probability for a Y-linked inversion subject to completely recessive mutations increases nearly linearly with $\mu$ and with the square root of $S$; the predicted absolute fixation probabilities with $\mu = 10^{-8}$ and $s = \{0.1, 0.25, 0.5\}$ are 0.0237, 0.0363, and 0.0506, respectively. These are also fairly close to the simulation values, but with an indication that the fixation probabilities are being overestimated for the larger $S$ values. With large $S$, although the series in Eq (A13) is absolutely convergent, the magnitude of successive terms may continue to increase for a long time, so that an evaluation of a large number of terms would be required for an accurate evaluation of $\tilde{Q}$.

#### Appendix B Jay et al. (2022)'s Computer Code

Jay et al. (2022)'s original computer code, downloaded from:
https://github.com/PaulYannJay/MutationShelteringTheory... (commit #c2b2f25),
is available as an online supplement to this article, and was archived on Zenodo on October 25, 2023 at:
https://zenodo.org/records/10041542.
A downloaded copy is also available on GitHub at:
https://github.com/colin-olito/Jay-2022-Comment.
The archived original simulation data used for plotting is available from:
https://figshare.com/authors/Paul_Jay/12493000.

The relevant lines of code for generating Figure 3c in Jay et al. (2022) appear on **L.30-58** of the file DeleteriousMutationSheltering_Figures_Simulations.R. The critical subsetting and summarizing steps where inversions lost by generation 20 are excluded from the data appear on **L.50-51**. Use of this subsetted data for plotting, and an additional step to exclude data for $s = 0$ appear on **L.58**:

```
30 ### Figure 3C ###
31 Simul=read.table(paste("~/Paper/ModelSexChrom/V3/CleanDataset/InversionTrajectories_N=1000_Fig3-S13-S15-S19.txt",sep=""), stringsAsFactors = F) #File containing all simulation with N=1000
32 colnames(Simul)=c("N", "u", "r", "h", "s", "Gen", "DebInv", "FinInv", "Rep",
33                   "MeanMutInv","MinMutInv","MaxMutInv","sdMutInv","FreqMutInv",
34                   "MeanMutNoInv","MinMutNoInv","MaxMutNoInv","sdMutNoInv","FreqMutNoInv",
35                   "InvFit", "NoInvFit","Freq","Chromosome")
36 Simul$Position="Y" #Create a position column
37 Simul[Simul$DebInv>10000000,]$Position="Autosome" #Inversion starting after position 10Mb are on the second chromosome
38 Simul[Simul$Position=="Y",]$Freq=Simul[Simul$Position=="Y",]$Freq * 4 # frequency of Y inversions in the population of Y chromosome, not the overall frequency
39 Simul$InvSize=Simul$FinInv - Simul$DebInv # Inversion size
40
41 Summary=Simul %>% group_by(N,u,r,h,s,InvSize,Position, Rep) %>% summarise(maxFreq=max(Freq), maxGen=max(Gen), InitMutNumb=min(MeanMutInv), MinSegMut=min(MeanMutNoInv)) # For each simulation,  grep its last generation (when the inversion was lost or fixed) and its maximum frequency
42 Summary$State="LostEarly" #Define an state defining inversion
43 Summary[Summary$maxGen>15020,]$State="LostLate" #Inversion lost after 20 generation or more
44 Summary[Summary$maxGen==24991,]$State="Segregating" #Inversion still segregating at simulation end
```

```
Summary[Summary$maxFreq>0.95,]$State="Fixed" #Inversion that reached above 0.95 frequency are considered fixed (for computation purpose, simulation stop when inversion fix,so sometime we do not observe inversion at 1.0)
Summary$StateCode=1 #For estimating the proportion of inversion fixed, note as 1 inversion fixed and 0 otherwise
Summary[Summary$State=="LostEarly",]$StateCode=0 #Define a code for each state
Summary[Summary$State=="LostLate",]$StateCode=0
Summary[Summary$State=="Segregating",]$StateCode=0
SumNoLostEarly=subset(Summary, (Summary$State!="LostEarly" & Summary$InitMutNumb>0)) #Remove inversions that were lost in fewer than 20 generations and those that are mutation free
DataSummary=SumNoLostEarly %>% group_by(N,u,r,h,s,InvSize, Position) %>% summarise(ProbSpread=mean(StateCode), MeanSegMut=mean(MinSegMut)) # For each set of parameter, compute the fraction of mutation fixed (only for not mutation-free inversion)

options(scipen=0) #non-scientific notation
Col=scales::viridis_pal(begin=0.0, end=0.8, option="A")(5) #Define color palette
n=2000000 #Focus on 2Mb inversions
DataSummary$u=factor(DataSummary$u, labels = c("mu==1␣%*%␣10^{-09}","mu==1␣%*%␣10^{-08}"), )

Base=ggplot(DataSummary[(DataSummary$InvSize == n & DataSummary$s<0),], aes( y=ProbSpread)) #Plot the result
```

Similar subsetting and summarizing steps are executed at the creation of each dataset used to generate plots of the fraction of fixed inversions, with data for $s = 0$ excluded using the plotting function or in a separate subsetting step:

```
### Figure S15 ### Same as figure 3C but with different inversion size
U="mu==1␣%*%␣10^{-08}"
DataSummary$InvSize=as.factor(paste0("Inversion␣size=",DataSummary$InvSize))
DataSummary$InvSize=relevel(DataSummary$InvSize,"Inversion␣size=500000")
Base=ggplot(DataSummary[(DataSummary$u == U & DataSummary$s<0),], aes( y=ProbSpread))
```

⋮

```
SumNoLostEarly10k=subset(Summary10k, (Summary10k$State!="LostEarly" & Summary10k$InitMutNumb>0))
DataSummary= SumNoLostEarly10k %>% group_by(N,u,r,h,s,InvSize, Position) %>% summarise(ProbSpread=mean(StateCode), MeanSegMut=mean(MinSegMut))
Col=scales::viridis_pal(begin=0.0, end=0.8, option="A")(5)
options(scipen=0)
```

```
DataSummary$InvSize=as.factor(paste0("Inversion␣size=",DataSummary$InvSize))
DataSummary$InvSize=relevel(DataSummary$InvSize,"Inversion␣size=500000")

Base=ggplot(DataSummary[(DataSummary$s<0),], aes( y=ProbSpread))
```

⋮

```
SumNoLostEarly=subset(summarySub_BLamb, (summarySub_BLamb$State!="LostEarly" & summarySub_BLamb$InitMutNumb>0))
DataSummary=SumNoLostEarly %>% group_by(N,u,r,s,InvSize, Position) %>% summarise(ProbSpread=mean(StateCode), MeanSegMut=mean(MinSegMut))

options(scipen=0)
Col=scales::viridis_pal(begin=0.2, end=0.6, option="A")(2)
options(scipen=0)
DataSummary$InvSize=as.factor(paste0("Inversion␣size=",DataSummary$InvSize))
DataSummary$InvSize=relevel(DataSummary$InvSize,"Inversion␣size=500000")
DataSub=DataSummary[(DataSummary$s<0),]
```

⋮

```
SumNoLostEarly=subset(Summary, Summary$State!="LostEarly") #Remove inversion that were lost in fewer than 20 generations
DataSummary=SumNoLostEarly %>% group_by(N,u,r,h,s,InvSize, FreqHaplo, Position) %>% summarise(ProbSpread=mean(StateCode)) # For each set of parameter, compute the fraction of mutation fixed (only for not mutation-free inversion)
```

### Appendix C  Supplementary Figures

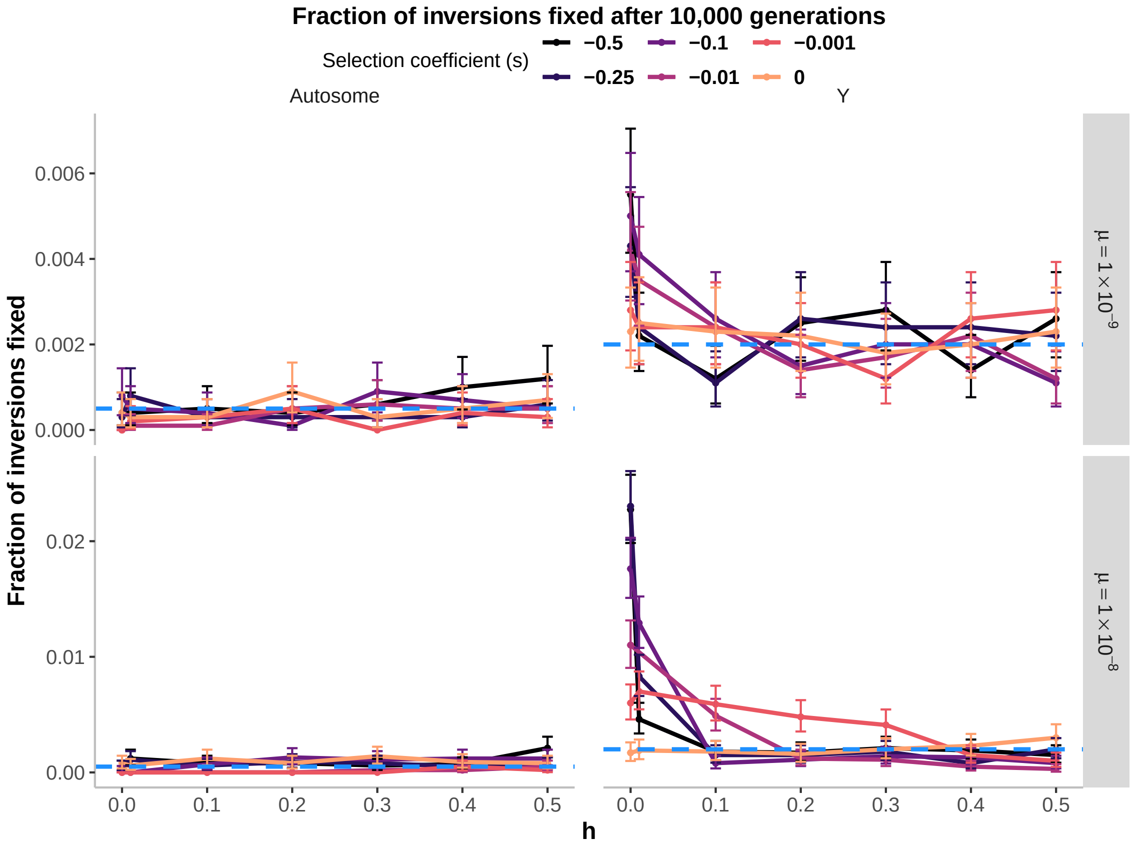

Figure S1: A slight modification of Fig. 2 from the main text (where the fraction of fixed inversions is calculated using Jay et al. (2022)'s full dataset), with figure panels arranged in the same way as Fig. 3c of (Jay et al. 2022), and Poisson 95% confidence intervals have been added to each estimate. This visualization of the same data clearly illustrates how closely the simulation results for $N = 1,000$ match the neutral expectations. Results are shown for dominance coefficients varying between completely recessive to additive fitness effects ($h = \{0.0, 0.01, 0.1, 0.2, 0.3, 0.4, 0.5\}$), selection coefficients ranging from quite weak to extremely strong selection ($s = \{0.0, 0.001, 0.01, 0.1, 0.25, 0.5\}$), and two per-base-pair mutation rates ($\mu = \{10^{-9}, 10^{-8}\}$). The expected proportion of fixations for a neutral gene first arising as a single copy is indicated by the blue horizontal dashed line (corresponding to $1/(2N)$ for an autosomal gene, and $2/N$ for a Y-linked gene.

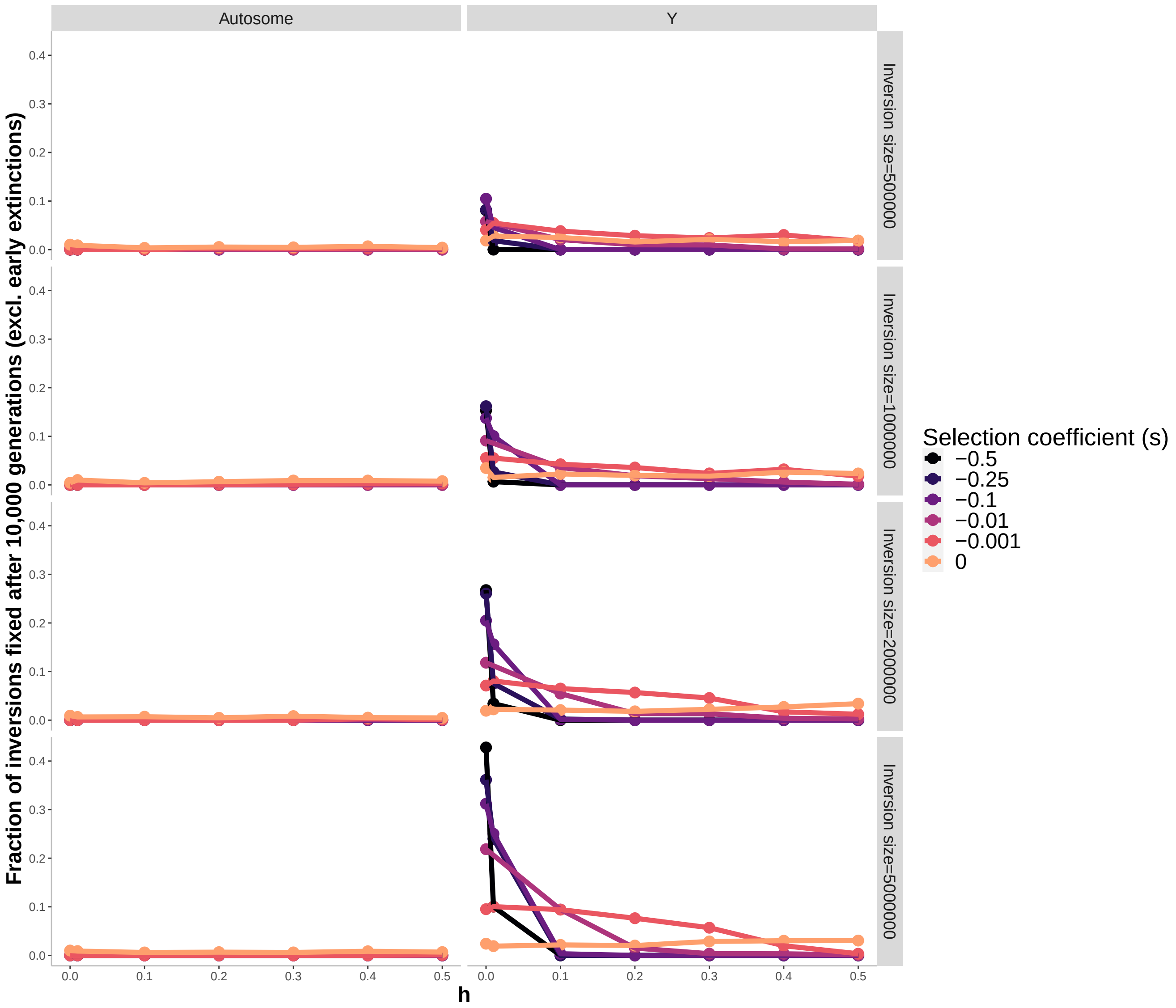

Figure S2: Reproduction of **Fig. S15** from Jay et al. (2022) using their data and code, but including data for $s = 0$, and with y-axis label and main figure title altered to reflect the correct interpretation of the data being plotted. The figure displays the frequency among 10,000 replicate simulations where independent single-copy inversion mutations go to fixation *conditioned on those inversions not going extinct in the first 20 generations, and not being mutation-free*, for a population size of $N = 1{,}000$, and a per-base-pair mutation rate of $\mu = 10^{-8}$. Results are shown for inversions spanning 500kb, 1Mb, 2Mb, and 5Mb of selected sites, dominance coefficients varying between completely recessive to additive fitness effects ($h = \{0.0, 0.01, 0.1, 0.2, 0.3, 0.4, 0.5\}$), and selection coefficients ranging from quite weak to extremely strong selection ($s = \{0.0, 0.001, 0.01, 0.1, 0.25, 0.5\}$).

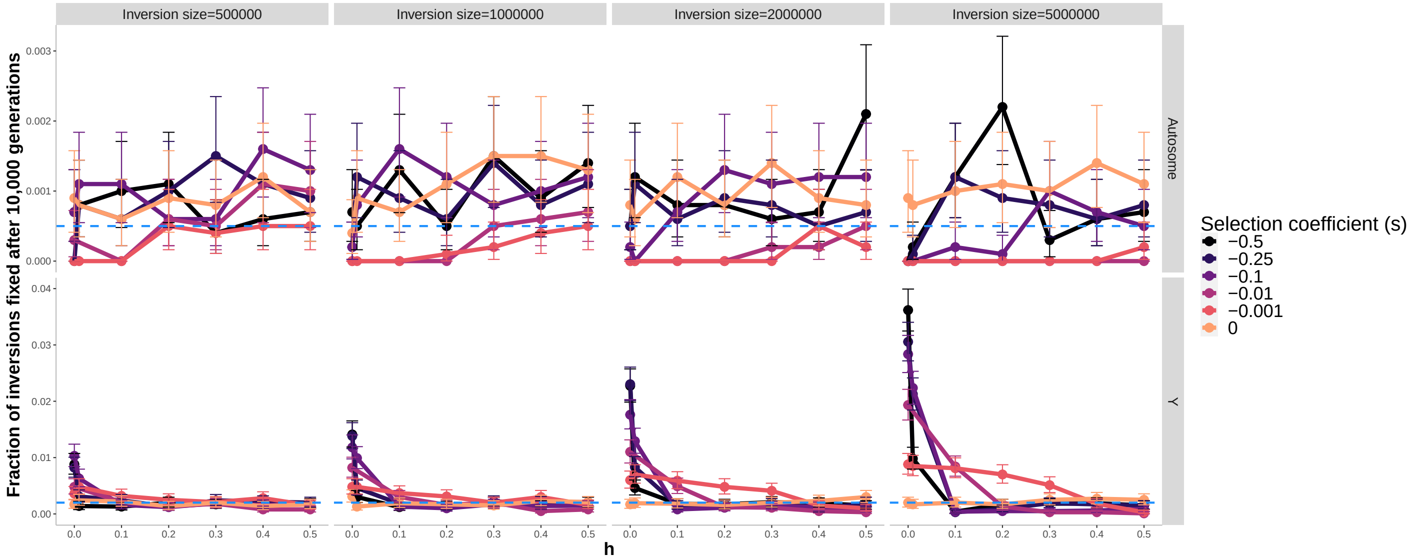

Figure S3: A revised presentation of the results from **Fig. S15** from Jay et al. (2022), where the fraction of fixed inversions among 10,000 replicate simulations is calculated using the full dataset, with figure panels rearranged so that autosomal and Y-linked inversions are plotted on different y-axis scales, and with Poisson 95% confidence intervals have been added to each estimate. These results can be interpreted as estimates of the overall probability of fixation for the inversions. Results are shown for inversions spanning 500kb, 1Mb, 2Mb, and 5Mb of selected sites for a population size of $N = 1,000$, and a per-base-pair mutation rate of $\mu = 10^{-8}$, with dominance coefficients varying between completely recessive to additive fitness effects ($h = \{0.0, 0.01, 0.1, 0.2, 0.3, 0.4, 0.5\}$), and selection coefficients ranging from quite weak to extremely strong selection ($s = \{0.0, 0.001, 0.01, 0.1, 0.25, 0.5\}$). The expected proportion of fixations for a neutral gene first arising as a single copy is indicated by the blue horizontal dashed line (corresponding to $1/(2N)$ for an autosomal gene, and $2/N$ for a Y-linked gene.

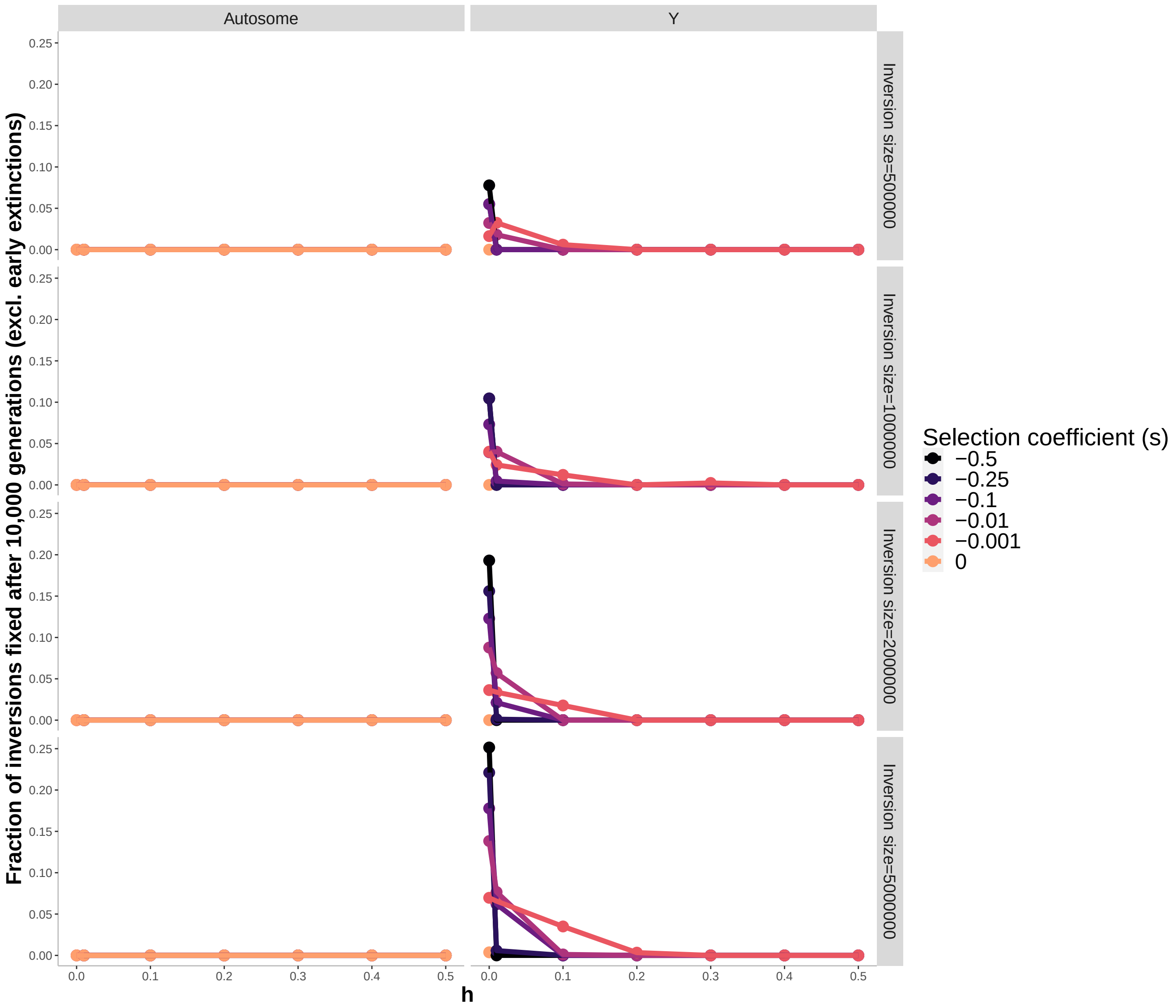

Figure S4: Reproduction of **Fig. S16** from Jay et al. (2022) using their data and code, but using the full dataset, and with y-axis label and main figure title changed to reflect the correct interpretation of the data being plotted. The figure displays the frequency among 10,000 replicate simulations where independent single-copy inversion mutations go to fixation *conditioned on those inversions not going extinct in the first 20 generations, and not being mutation-free*, for a population size of $N = 10,000$, and a per-base-pair mutation rate of $\mu = 10^{-8}$. Results are shown for inversions spanning 500kb, 1Mb, 2Mb, and 5Mb of selected sites, with dominance coefficients varying between completely recessive to additive fitness effects ($h = \{0.0, 0.01, 0.1, 0.2, 0.3, 0.4, 0.5\}$), and selection coefficients ranging from quite weak to extremely strong selection ($s = \{0.0, 0.001, 0.01, 0.1, 0.25, 0.5\}$).

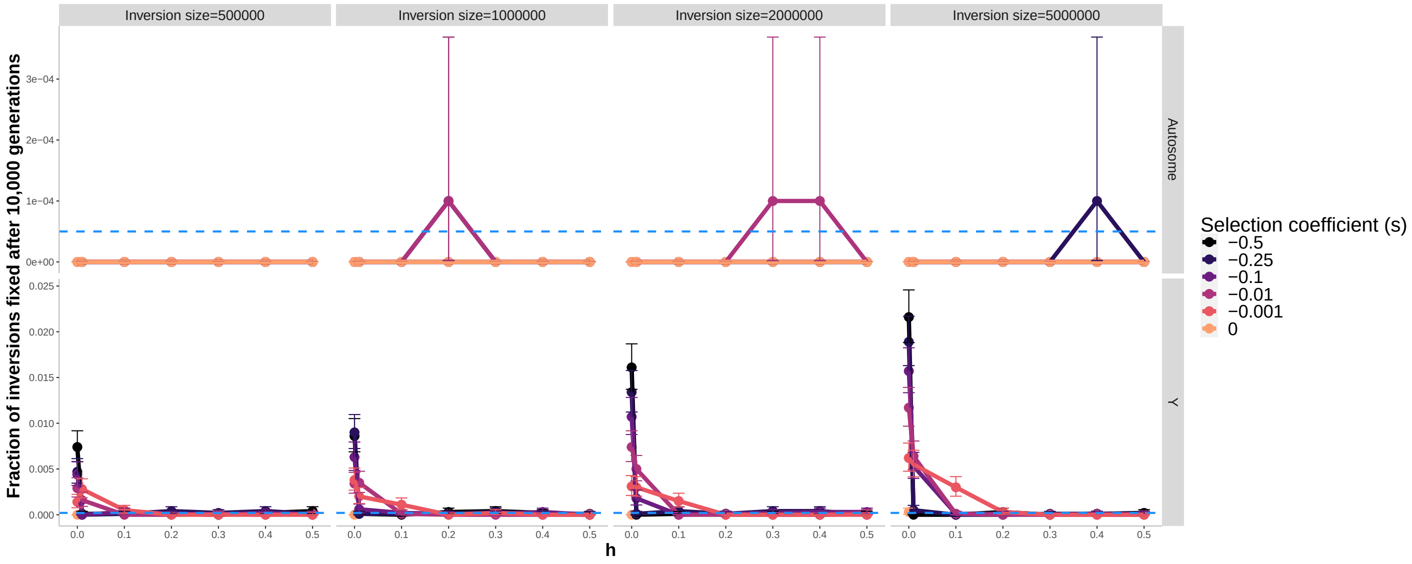

Figure S5: A revised presentation of the results from **Fig. S16** from Jay et al. (2022), where the fraction of fixed inversions among $10,000$ replicate simulations is calculated using the full dataset, the figure panels have been rearranged so that autosomal and Y-linked inversions are plotted on different y-axis scales, and Poisson 95% confidence intervals have been added to each estimate. These results can be interpreted as estimates of the overall probability of fixation for the inversions. Results are shown for inversions spanning 500kb, 1Mb, 2Mb, and 5Mb of selected sites for a population size of $N = 10,000$, a per-base-pair mutation rate of $\mu = 10^{-8}$, and with dominance coefficients varying between completely recessive to additive fitness effects ($h = \{0.0, 0.01, 0.1, 0.2, 0.3, 0.4, 0.5\}$), and selection coefficients ranging from quite weak to extremely strong selection ($s = \{0.0, 0.001, 0.01, 0.1, 0.25, 0.5\}$). The expected proportion of fixations for a neutral gene first arising as a single copy is indicated by the blue horizontal dashed line (corresponding to $1/(2N)$ for an autosomal gene, and $2/N$ for a Y-linked gene. *Note: There appear to be missing data for Y-linked inversions with $s = 0$ in the origina dataset provided by Jay et al. (2022).*

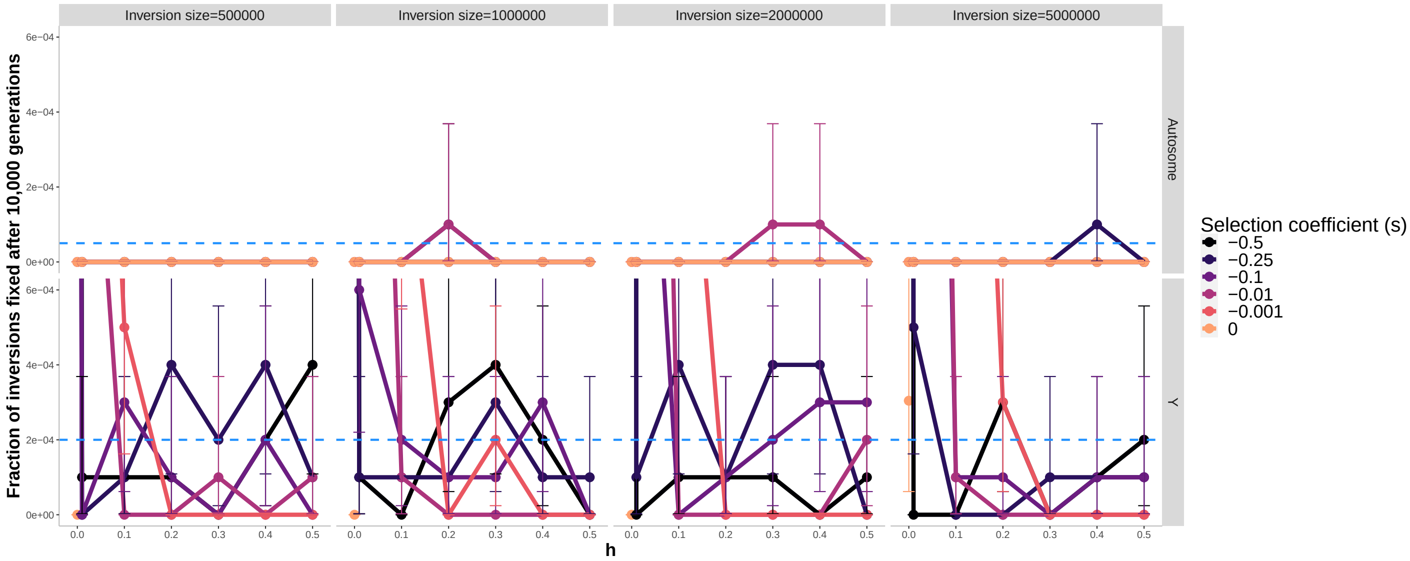

Figure S6: Same as fig. S5, but zoomed in to a maximum y-axis value of $6/N$ to more clearly visualize results for $h > 0.01$. The plot still shows the fraction of fixed inversions among $10,000$ replicate simulations calculated using the full dataset, and with figure panels rearranged with Poisson 95% confidence intervals for each estimate. These results can be interpreted as estimates of the overall probability of fixation for the inversions. Results are shown for inversions spanning 500kb, 1Mb, 2Mb, and 5Mb of selected sites for a population size of $N = 10,000$, a per-base-pair mutation rate of $\mu = 10^{-8}$, and with dominance coefficients varying between completely recessive to additive fitness effects ($h = \{0.0, 0.01, 0.1, 0.2, 0.3, 0.4, 0.5\}$), and selection coefficients ranging from quite weak to extremely strong selection ($s = \{0.0, 0.001, 0.01, 0.1, 0.25, 0.5\}$). The expected proportion of fixations for a neutral gene first arising as a single copy is indicated by the blue horizontal dashed line (corresponding to $1/(2N)$ for an autosomal gene, and $2/N$ for a Y-linked gene. *Note: Here, it is more clear that there is missing data for Y-linked inversions with $s = 0$ in the origina dataset provided by Jay et al. (2022).*

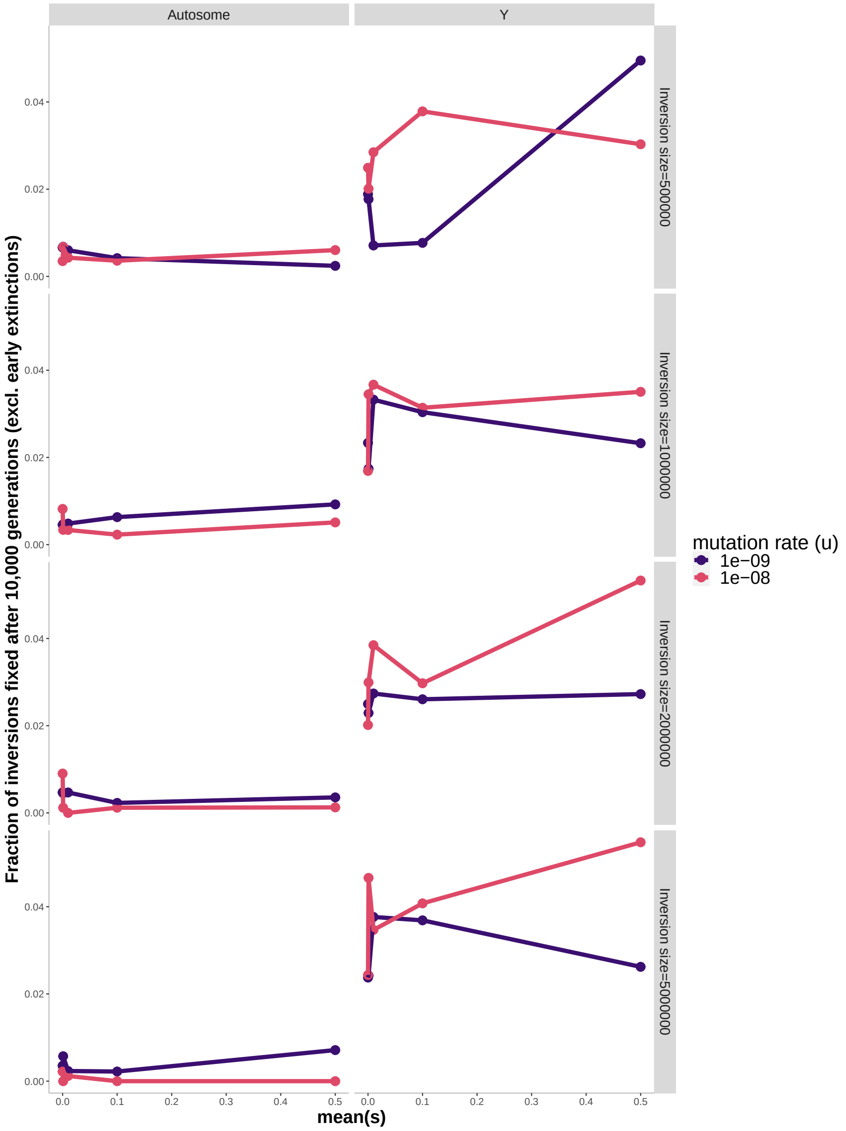

Figure S7: Reproduction of the original **Fig. S17** from Jay et al. (2022) using their data and code, including data for $s = 0$, and with y-axis label and main figure title altered to reflect the correct interpretation of the data being plotted. The figure displays the frequency among 10,000 replicate simulations where independent single-copy inversion mutations go to fixation *conditioned on those inversions not going extinct in the first 20 generations, and not being mutation-free*, for a population size of $N = 1,000$, a per-base-pair mutation rate of $\mu = 10^{-8}$, but with deleterious mutationsthat have their fitness effects drawn from a gamma distribution with a shape of 0.2, and their dominance coefficient $h$ randomly sampled from the possible values of $0, 0.001, 0.01, 0.1, 0.25, 0.5$ with uniform probabilities. Results are shown for inversions spanning 500kb, 1Mb, 2Mb, and 5Mb of selected sites, with the mean of the selection coefficient values (mean of the gamma distribution) of $\bar{s} = \{0.0, 0.001, 0.01, 0.1, 0.5\}$. *Note: To include data for* $s = 0$*, we changed the scaling of the x-axis so that it is not on a* $log_{10}$ *scale.*

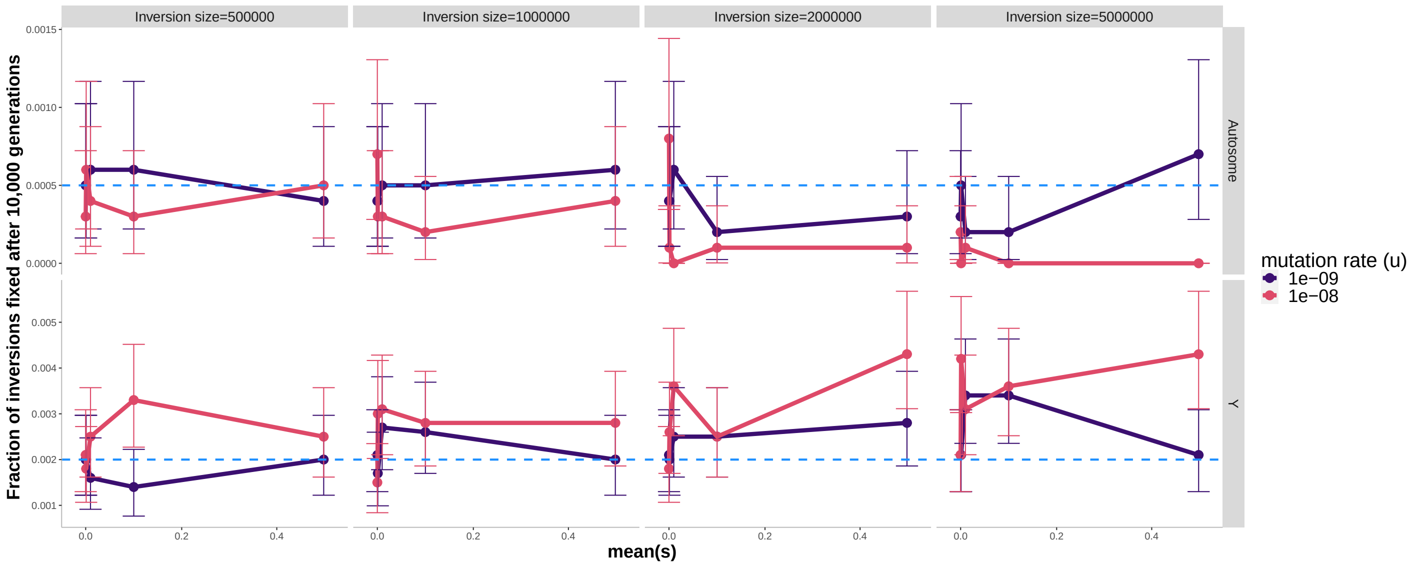

Figure S8: A revised presentation of the results from **Fig. S17** from Jay et al. (2022), where the fraction of fixed inversions among 10,000 replicate simulations is calculated using the full dataset, with figure panels rearranged so that autosomal and Y-linked inversions are plotted on different y-axis scales, and with Poisson 95% confidence intervals added to each estimate. These results can be interpreted as estimates of the overall probability of fixation for the inversions. for a population size of $N = 1,000$, a per-base-pair mutation rate of $\mu = 10^{-8}$, but with deleterious mutations that have their fitness effects drawn from a gamma distribution with a shape of 0.2, and their dominance coefficient $h$ randomly sampled from the possible values of $0, 0.001, 0.01, 0.1, 0.25, 0.5$ with uniform probabilities. Results are shown for inversions spanning 500kb, 1Mb, 2Mb, and 5Mb of selected sites, with the mean of the selection coefficient values (mean of the gamma distribution) of $\bar{s} = \{0.0, 0.001, 0.01, 0.1, 0.5\}$. The expected proportion of fixations for a neutral gene first arising as a single copy is indicated by the blue horizontal dashed line (corresponding to $1/(2N)$ for an autosomal gene, and $2/N$ for a Y-linked gene. *Note: To include data for $s = 0$, we changed the scaling of the x-axis so that it is not on a $\log_{10}$ scale.*
